# Transcriptome Analysis of Chloride Intracellular Channel Knockdown in *Drosophila* Identifies Oxidation-Reduction Function as Possible Mechanism of Altered Sensitivity to Ethanol Sedation

**DOI:** 10.1101/2021.01.20.427413

**Authors:** Rory M. Weston, Rebecca E. Schmitt, Mike Grotewiel, Michael F. Miles

## Abstract

Chloride intracellular channels (CLICs) are a unique family of evolutionarily conserved metamorphic proteins, switching between stable conformations based on redox conditions. CLICs have been implicated in a wide variety biological processes including ion channel activity, apoptosis, membrane trafficking, and enzymatic oxidoreductase activity. Understanding the molecular mechanisms by which CLICs engage in these activities is an area of active research. Here, the sole *Drosophila melanogaster* ortholog, *Clic*, was targeted for RNAi knockdown to identify genes and biological processes associated with *Clic* expression. *Clic* knockdown had a substantial impact on global transcription, altering expression of over 9% of transcribed *Drosophila* genes. Overrepresentation analysis of differentially expressed genes identified enrichment of 23 Gene Ontology terms including Cytoplasmic Translation, Oxidation-Reduction Process, Heme Binding, Membrane, Cell Junction, and Nucleolus. The top term, Cytoplasmic Translation, was enriched almost exclusively with downregulated genes. Drosophila *Clic* and vertebrate ortholog *Clic4* have previously been tied to ethanol sensitivity and ethanol-regulated expression. *Clic* knockdown-responsive genes from the present study were found to overlap significantly with gene sets from 4 independently published studies related to ethanol exposure and sensitivity in *Drosophila*. Bioinformatic analysis of genes shared between these studies revealed an enrichment of genes related to amino acid metabolism, protein processing, oxidation-reduction processes, and lipid particles among others. To determine whether the modulation of ethanol sensitivity by *Clic* may be related to co-regulated oxidation-reduction processes, we evaluated the effect of hyperoxia on ethanol sedation in *Clic* knockdown flies. Consistent with previous findings, *Clic* knockdown reduced acute ethanol sedation sensitivity in flies housed under nomoxia. However, this effect was reversed by exposure to hyperoxia, suggesting a common set of molecular-genetic mechanism may modulate each of these processes. This study suggests that *Drosophila Clic* has a major influence on regulation of oxidative stress signaling and that this function overlaps with the molecular mechanisms of acute ethanol sensitivity in the fly.

## Introduction

Chloride intracellular channels (CLICs) are a family of evolutionarily conserved proteins with unique metamorphic properties and a host of highly diverse, yet poorly understood biological functions. Vertebrates possess 6 highly similar chloride intracellular channel paralogs and orthologs are also found in invertebrates including *Caenorhabditis elegans* and *Drosophila melanogaster* (1). The biological functions of CLICs have been difficult to ascertain, but insight has been gained through knockout models in mice and *C. elegans*. Although viable, animals deficient for CLICs exhibit a diverse array of phenotypes including defective excretory canal formation in *C. elegans* (2) and impaired angiogenesis (3,4), and wound healing in mice (5). Work in knockout models has been complemented by *in vitro* studies and the overall list of functions associated with CLICs now includes roles in ion channel activity (6–8), membrane trafficking (9,10), apoptosis (11,12), TGF-beta signaling (5,13,14), tubulogenesis (2,3,9), innate immunity (15,16), and oxidoreductase enzymatic activity (17) among others. Unfortunately, little progress has been made in identifying the molecular mechanisms by which CLICs engage in these diverse biological processes and much remains to be elucidated.

As members of a rare class of metamorphic proteins, CLICs can alter their three-dimensional structure in a ligand-free environment in response to changes in redox conditions (7,18,19). Under oxidizing conditions, CLICs can rearrange their tertiary structure and spontaneously insert into membranes where they demonstrate an ability to conduct ions across membranes through an unknown mechanism (6–8). The selectivity of CLICs for anions, let alone chloride, has been challenged suggesting the channels may better resemble membrane pores (20). Under reducing conditions, CLICs tend towards a soluble globular conformation which has been associated with enzymatic oxidoreductase activity *in vitro* (17). This finding is not entirely surprising considering the structural homology of CLICs and omega class glutathione S-transferase (GST) enzymes (6,21). General features of CLICs such as their resemblance to omega class GSTs, ability to interconvert structures and conduct ions across membranes are largely conserved between vertebrates to invertebrates (22). One major distinction between invertebrate and vertebrate CLICs is the presence of a two-cysteine redox active site, which is disrupted in *C. elegans* paralogs *exl-1* and *exc-4*, but maintained in the sole *Drosophila* ortholog, *Clic*. This active site has been linked to binding of CLICs to lipid bilayers after oxidation, which is true of vertebrate and *Drosophila* CLICs, but not *C. elegans* (22). This active site motif may also be necessary for glutathione binding and oxidoreductase enzymatic activity (17).

Growing evidence has linked CLICs to ethanol-related behaviors and identified them as a potentially important risk factor for alcohol use disorder (AUD) in humans. Expression of chloride intracellular channel 4 (*Clic4*) is downregulated in the brains of postmortem human alcoholics (23) and part of an ethanol-responsive gene network in mouse brain (24). *Clic4* has been shown to be induced in mouse brain by acute ethanol (25,26) and overexpression of *Clic4* decreased sensitivity to ethanol sedation in mice (25). In the same study, transposon disruption of *Drosophila Clic* and mutation of *C. elegans exc-4* were also shown to decrease ethanol sedation sensitivity. In a separate study, RNAi knockdown of *Drosophila Clic* replicated these findings by reducing sensitivity to ethanol sedation (27). These findings are significant considering the possible role of low initial ethanol sensitivity as a risk factor in the development of AUD in humans (28,29). Similar to many other biological functions associated with CLICs, the molecular mechanisms by which they alter ethanol sensitivity is presently unknown.

The present study has taken steps to address these gaps in understanding the molecular mechanisms of CLIC action and role in ethanol behaviors by using the power of *Drosophila* genetics to knock-down *Clic* expression selectively in neurons and characterizing the consequent transcriptomic response. Investigation of transcriptome networks resulting from *Clic* knockdown would not only add to our knowledge on *Clic* function, but might also increase our understanding of the neurobiology underlying ethanol sedation sensitivity in the fly. Our findings provide validation for published roles for CLICs, identify potentially novel functions and genetic interactions that shed light on the nature of chloride intracellular channel biology, and show a remarkable conservation of transcriptome responses to *Clic* knockdown, genes involved in oxidative stress and molecular mechanisms relating to ethanol sedation sensitivity in *Drosophila*.

## Materials and Methods

### *Drosophila* Husbandry, Genetics, and Behavioral Studies

Flies harboring the neuron-selective *elav*-Gal4 driver and/or *Clic* UAS-RNAi transgenev105975 were reared, crossed, and evaluated for sensitivity to sedation to vapor from 85% ethanol as previously described (27). Flies were placed in sealed plastic containers containing 95% O_2_ (charged twice daily) for exposure to hyperoxia. Survival following repeated hyperoxia exposures was evaluated as previously described (30).

### RNA Extraction and Microarray Preparation

RNA was extracted from fly heads as previously described (30). Microarray preparation performed per standard Affymetrix protocol using GeneChip *Drosophila* Gene 1.0 ST arrays (ThermoFisher Scientific #902155). Hybridization, washing, and scanning performed per manufacturer specifications by VCU Massey Cancer Center Tissue and Data Acquisition and Analysis Core.

### Microarray Analysis

All microarray data processing, statistical analysis, and bioinformatics were performed in R v3.5.1 (31) using R Studio v1.1.456 (32) unless otherwise stated. Microarray CEL files were preprocessed with the R package Oligo v1.44.0 (33) for quality control visualization and background subtraction and normalization was performed with the default robust multi-array average (RMA) method. Release 36 of the corresponding Affymetrix *Drosophila* Gene 1.0 ST array transcript annotations were used. Differential gene expression analysis was performed with the R package Limma v3.36.5 (34) using gene-level linear model fitting and empirical Bayesian smoothing of standard errors per the default workflow. P-values were adjusted using the false discovery rate method (35) and a cutoff of less than or equal to 0.05 was applied for significant differential expression. Plotting for these analyses was performed with the R package ggplot2 v3.0.0 (36). Principal component analysis (PCA) plotting performed by ggbiplot R package v0.55 (37) with computed normal confidence ellipses feature enabled. Microarray data files have been deposited at the Gene Expression Omnibus under accession number GSE164090 (GEO, https://www.ncbi.nlm.nih.gov/geo/).

### Bioinformatics

Functional enrichment analysis of differentially expressed genes found with Limma analysis was performed using the web-based tool DAVID (https://david.ncifcrf.gov/) (38). Databases examined included the Kyoto Encyclopedia of Genes and Genomes (KEGG) (39,40) and Gene Ontology (GO) categories of Biological Processes, Cellular Components, and Molecular Functions (39,41). A p-value cutoff of 0.01 was applied to all GO terms and terms with > 90% redundancy were removed. Significantly enriched terms were visually explored using the R package GOplot v.1.0.2 (42) to produce the representative plots in Fig 3. The web-based tool GeneWeaver (https://geneweaver.org/) was used to perform an integrative genomic analysis across multiple published *Drosophila* gene sets (43). Using the HiSim Graph tool, differentially expressed genes from the present *Clic* knockdown were found to have significant Jaccard similarity with four published *Drosophila* ethanol exposure (44–46) and sedation sensitivity (47) gene sets (GS137794, GS75550, GS137795, and GS75562 respectively). These four gene sets were combined to create a union set of ethanol-sensitive genes, which was then compared to the *Clic* knockdown-altered genes using a Fisher’s exact test-based method provided in the R package GeneOverlap v.1.16.0 (48). Genes found to overlap between the ethanol-sensitive union and *Clic* knockdown sets were submitted for bioinformatic analysis by DAVID in order to identify enriched functional terms common between ethanol and *Clic* knockdown-sensitive genes.

The DRSC Integrative Ortholog Prediction Tool (https://www.flyrnai.org/cgi-bin/DRSC_orthologs.pl) was used to obtain human orthologs for the *Clic* knockdown differentially expressed gene list (49). In cases where multiple orthologs were found for a single *Drosophila* gene, only the top ortholog according to parameters *w_score, best_rev, sim_score*, and *identity* was used. The top 150 up and downregulated orthologs were then provided to the CLUE web-based tool for Connectivity Map (CMap) analysis (https://clue.io/), which compares the input transcriptomic signature with that of 476,251 transcriptomic signatures obtained from in vitro exposure of 9 human cell lines to 27,927 distinct chemical or RNAi perturbagens (50). Only perturbagen signatures having connectivity scores (tau) > 90 or <-90 are reported here.

## Results

### Differential Gene Expression Following *Clic* Knockdown

A neuron-specific Gal4 expressing *Drosophila* strain (*elav*-Gal4) was crossed to a UAS-dependent *Clic*-targeting RNAi strain (v105975/+), producing a neuronally-selective *Clic* knockdown strain (*elav*/v105975, Fig 1). To identify genes dysregulated by *Clic* knockdown, total RNA was extracted from fly heads for each strain and analyzed using Affymetrix Genome 2.0 Arrays, which quantifies expression of more than 18,500 Drosophila transcripts. Principal component analysis (PCA) of robust multi-array average (RMA) corrected probeset intensities revealed clear separation of the *elav*/v105975 knockdown and *elav*/+ control fly strain samples (Fig 2a).

**Fig 1.**
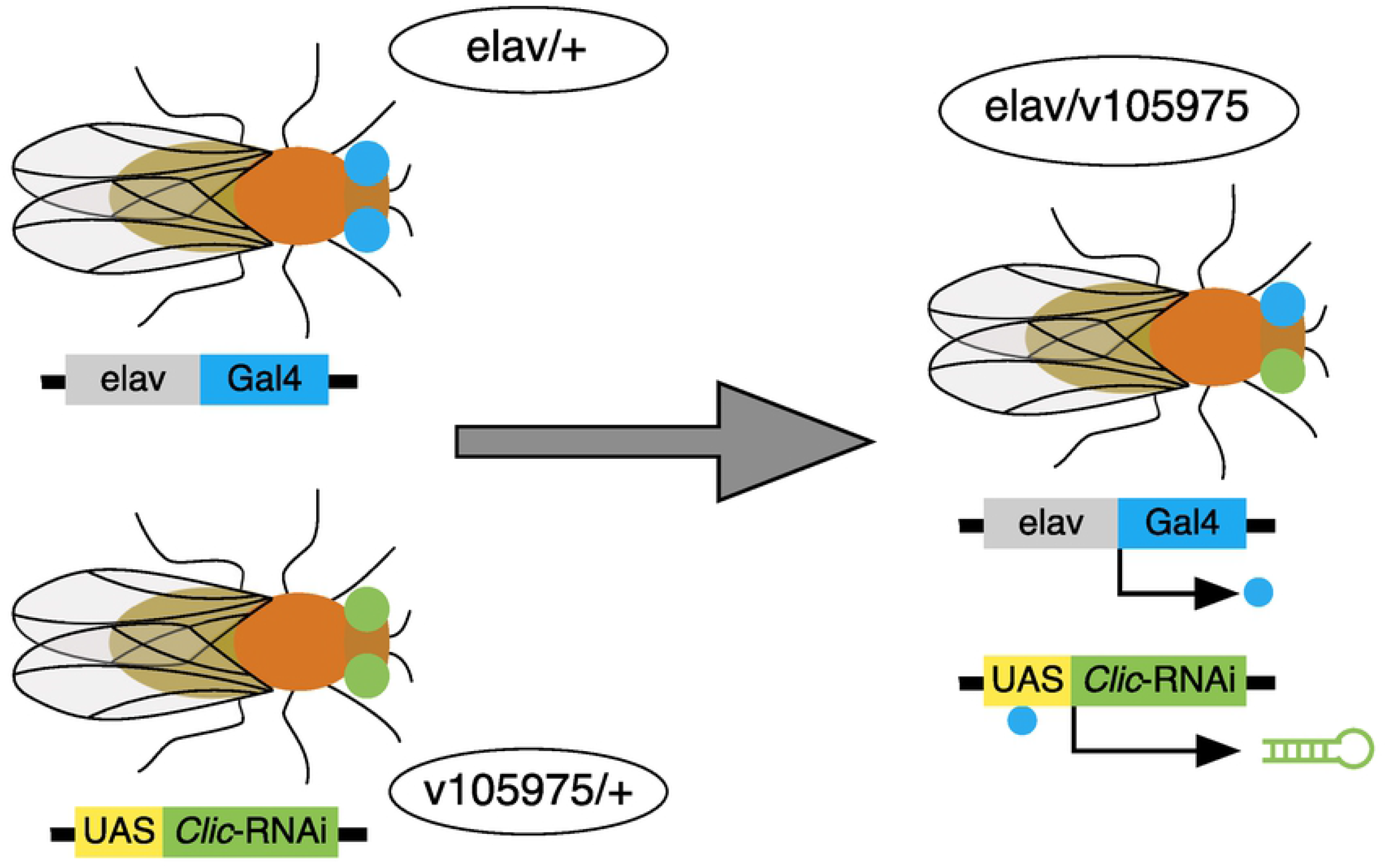
Overview of *Clic* knockdown approach. Schematic depicting breeding scheme for neuronal-specific Gal4 expression under the *elav* promoter driving UAS activated *Clic*-RNAi expression in *Drosophila*.

**Fig 2.**
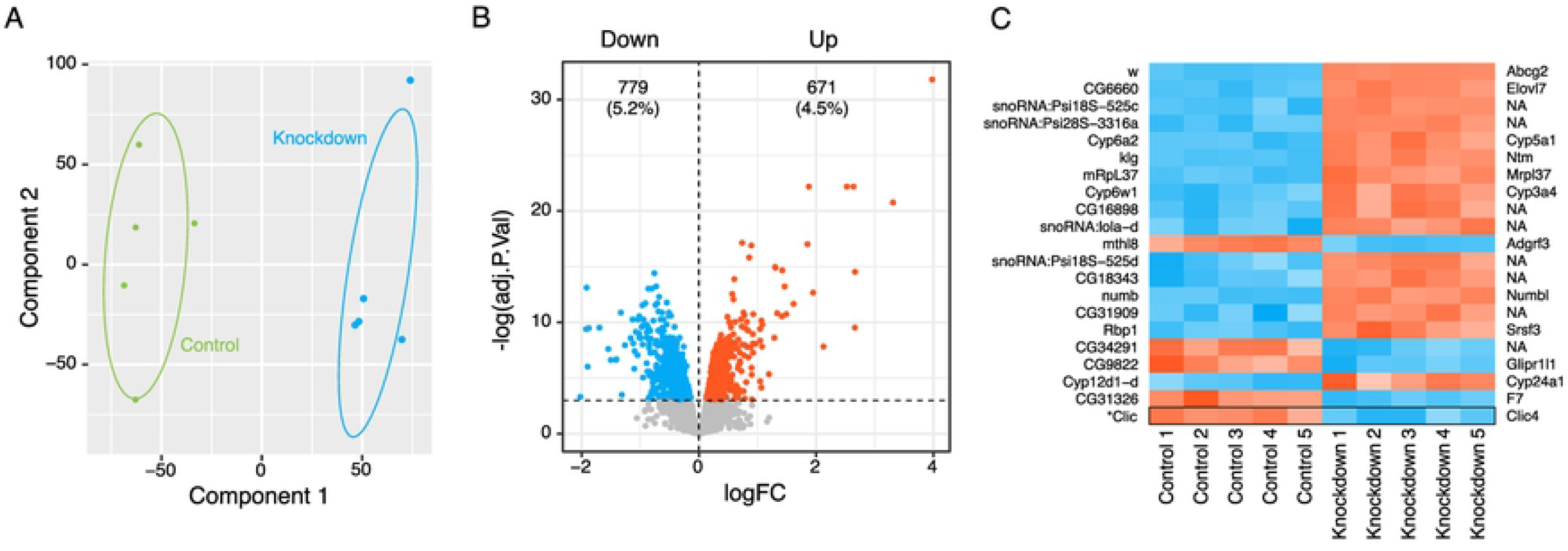
*Clic* knockdown-responsive gene expression. (a) PCA plot depicting expression profiles for control (*elav*/+) and *Clic* knockdown flies (*elav*/v105975) with normal confidence ellipses. (b) Volcano plot for complete differential gene expression results, highlighting significantly downregulated (blue) and upregulated (red) genes (FDR < 0.05). (c) Heatmap of top 20 differentially regulated genes, ranked by FDR. Fly genes are listed on left and corresponding human orthologs on right (NA indicates no clear ortholog). *Clic* expression added to bottom row of heatmap for clarity.

Differential gene expression analysis of the two strains identified 1,450 differentially expressed genes after applying a false discovery rate (FDR) cutoff of 0.05 (Fig 2b, S1 Table). Differentially expressed genes represented 9.7% of the total genes assessed, and although split fairly evenly, showed a trend towards overall downregulation. Human orthologs for the top 20 differentially expressed genes according to FDR include multiple cytochrome p450 enzymes (Cyp) as well as examples of membrane-bound (Abcg2, Elovl7, Ntm, and Glipr1l1) and translation-associated (Mrpl37 and Srsf3) proteins (Fig 2c). The knockdown strain (*elav*/v105975) had twice the number of copies of selectable marker gene mini-white (w) as the control strain (*elav*/+), rendering it the top differentially expressed gene as expected. The knockdown target gene, *Clic*, was expressed at 59% of *elav*/+ control fly levels, confirming previously reported knockdown using the same UAS-RNAi strategy measured by real-time PCR (27).

To explore the possibility of RNAi expression leakage in the Gal4-UAS system, v105975/+ RNAi-only controls were assessed alongside the *elav*/v105975 knockdown and *elav*/+ Gal4-only control strains during differential gene expression analysis. v105975/+ flies showed a 15% reduction in *Clic* expression compared to *elav*/+ controls, suggesting expression of RNAi molecules is occurring in the absence of a Gal4 driver in v105975/+ animals (S1 Table). While the knockdown magnitude in v105975/+ flies was much less than in the *elav*/v105975 knockdown strain, it did result in substantial differential gene expression (S1 Fig, panel a). However, only 54 genes were differentially expressed between the v105975 RNAi-only control and *elav*/v105975 knockdown strain and all but 14 of those were also differentially expressed between the *elav*/v105975 knockdown and *elav*/+ control strains (S1 Fig, panel b). Considering this high degree of similarity, the v105975 RNAi-only genotype was effectively a lower dose knockdown and was therefore omitted from the rest of the bioinformatic analyses in order to focus on the full *elav*/v105975 knockdown.

### Perturbed Oxidation-Reduction and Cytoplasmic Translation

To objectively screen the large list of differentially expressed genes for meaningful biological patterns, functional over-representation analysis was performed using the GO classification system. Twenty-three non-redundant GO terms with p-values < 0.01 were identified from all three GO categories (Biological Processes, Molecular Functions, & Cellular Components) and reflected trends observed in the top 20 differentially expressed genes (Fig 3a, S2 Table). The top 6 overrepresented GO terms according to p-value included Biological Processes Cytoplasmic Translation and Oxidation-Reduction Process, Molecular Functions Heme Binding, Cellular Components Membrane, Cell Junction, and Nucleolus (Fig 3a-d). Differentially expressed genes localized to the nucleolus and those involved in cytoplasmic translation, oxidation-reduction processes, and heme binding are largely downregulated whereas those localized to membranes or cell junctions are mostly upregulated (Fig 3a-c). Despite having large z-scores for overall direction of regulation (Fig 3a,b), terms such as Oxidation-Reduction Process and Cell Junction possessed examples of genes with opposing directions of regulation, highlighting the complex but specific molecular responses to *Clic* knockdown (Fig 3c). For example, Cyp genes were particularly overrepresented among top *Clic* knockdown-responsive genes, but showed considerable variation in direction of regulation, despite a low overall z-score for their parent term Oxidation-Reduction Process.

**Fig 3.**
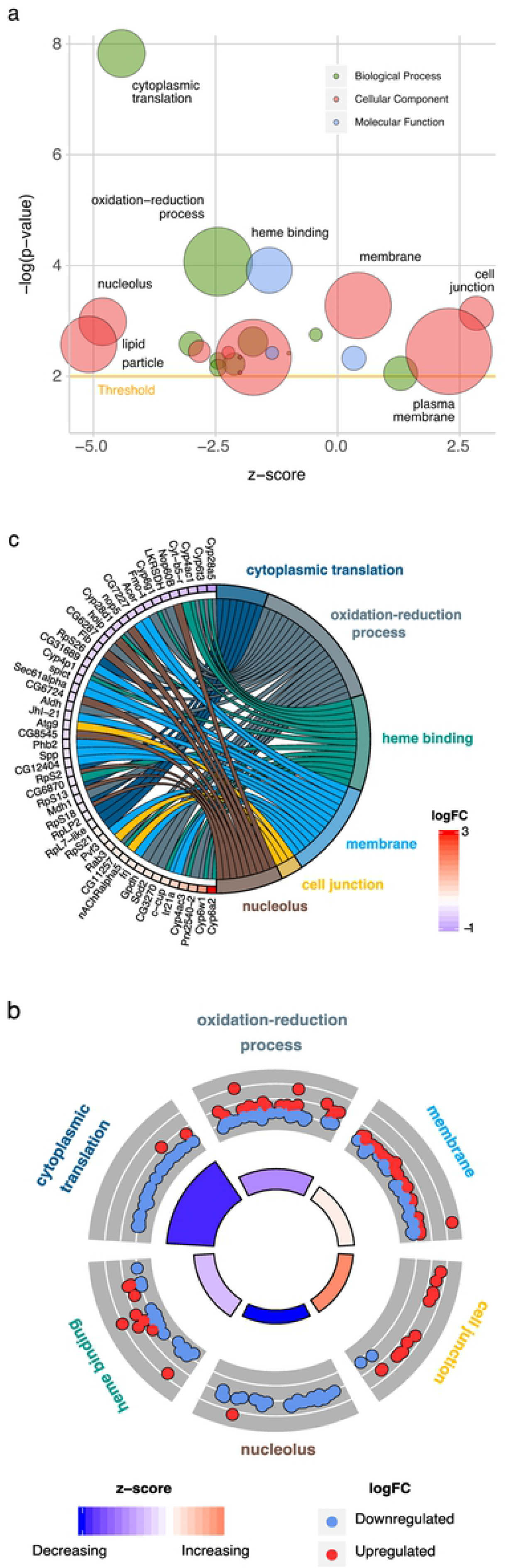
GO Terms Enriched by *Clic* Knockdown. (a) GO terms significantly affected by *Clic* knockdown with a p-value cutoff set to 0.01. Bubble radius is proportionate to term size in total number of genes and z-score represents overall direction of regulation of differentially expressed genes. (b) Circle plot depicting top 6 GO terms according to enrichment p-value. Outer ring corresponds to regulation of individual genes (logFC) within a term while inner ring corresponds to term enrichment p-value (bar height) and direction of regulation z-score (color). (c) Top 6 GO terms and top 50 differentially regulated genes from union of all 6 terms’ gene sets, depicted by gene name.

### Overlap with Ethanol-Regulated Genes

To gain further insight into the biological functions associated with *Clic*, the knockdown gene expression profile was screened against the large database of other transcriptomic studies available through GeneWeaver (Baker 2012). The most similar gene sets identified, having significant Jaccard Index scores (p < 0.05), were obtained from 4 transcriptomic studies related to ethanol exposure (44–46) and sedation sensitivity (47) in *Drosophila* (Fig 4a). A union of these ethanol-responsive gene sets was intersected with the *Clic* knockdown-responsive gene set and a significant overlap of 366 genes (p = 1.8×10−29, OR = 2.2) was found (Fig 4b, **S3 Table**). These genes were overrepresented in multiple GO terms and KEGG pathways, including metabolic and redox processes, sensory perception, protein processing, and transport among others (Fig 4c).

**Fig 4.**
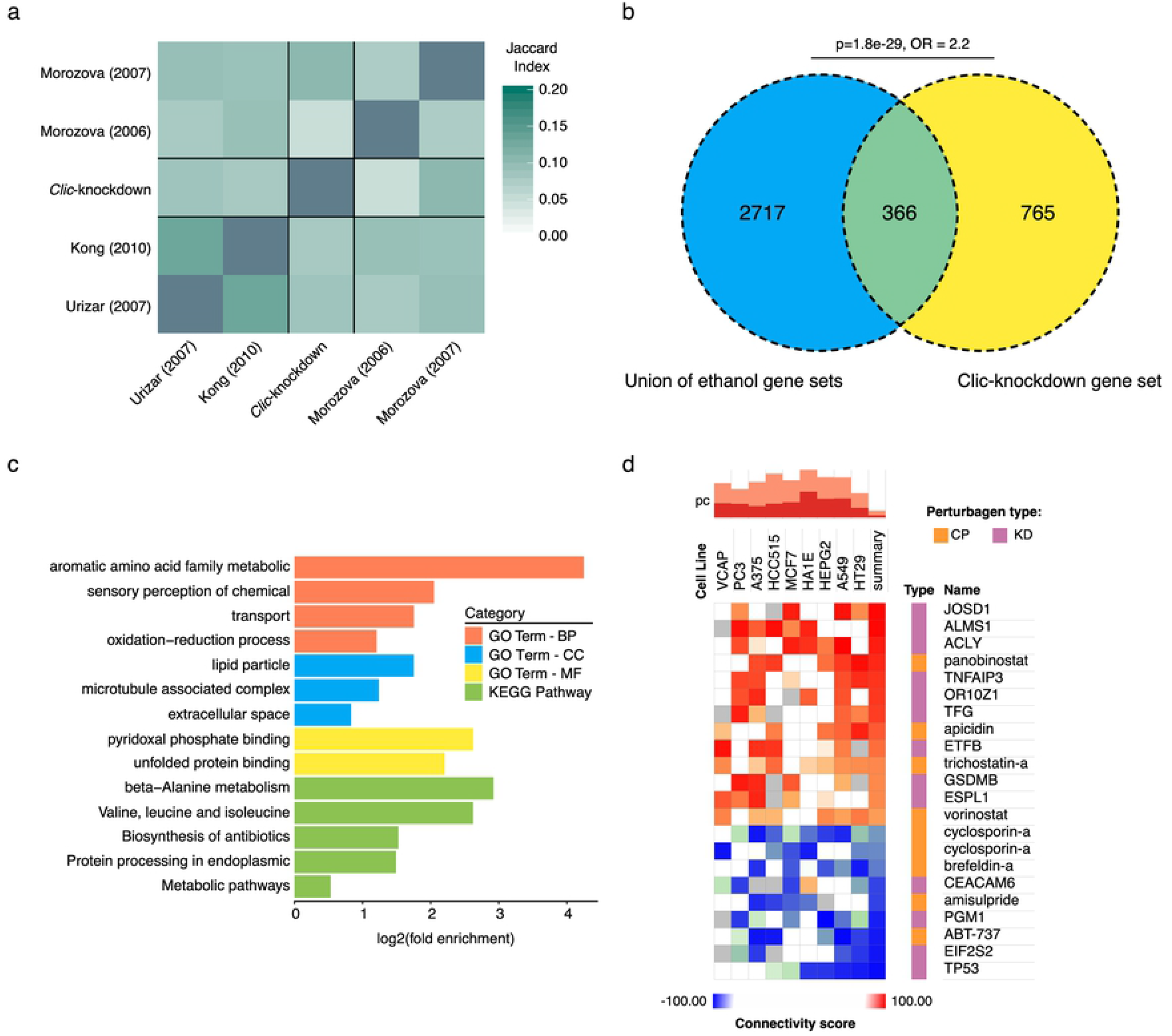
Gene Sets Overlapping with *Clic* Knockdown. (a) Heatmap showing Jaccard similarity between the *Clic* knockdown-sensitive gene set and 4 *Drosophila* ethanol-related gene sets obtained through GeneWeaver. Genes shared between the union of the 4 ethanol-related gene sets and the *Clic* knockdown-responsive gene set shown in (b) along with their GO functional enrichment analysis (c). (d) CMap analysis of perturbagen transcriptomic signatures with high positive (red, tau > 90) and negative (blue, tau < 90) connectivity with the *Clic* knockdown transcriptomic signature among 9 human cell lines. Assayed perturbagens include compounds (CP) and gene knockdowns (KD).

How *Clic* modulates resistance to ethanol sedation is not known and as a member of a class of proteins with incompletely characterized function, identification of selective pharmacological activators and inhibitors for more direct investigation is challenging. Using the cloud-based CLUE platform for CMap analysis, the transcriptomic signature of *Clic* knockdown was correlated with transcriptomic signatures of over 19,000 small molecules previously tested in human cell lines. This approach was an attempt to produce a list of small molecules with transcriptomic signatures positively or negatively connected to the signature of *Clic* knockdown, thereby identifying potentially novel pharmacological modulators of *Clic* function. The CMap screen was able to identify 22 perturbagens, either chemical small molecules or RNAi, that showed significant connectivity (tau > 90 or < −90) with transcriptomic signature of *Clic* knockdown (Fig 4d). Among chemical perturbagens, *Clic* knockdown was positively connected with histone deacetylase inhibitors (HDI) apicidin, panobinostat, trichostatin-a, and vorinostat and negatively connected to immunosuppressant cyclosporin-a, unfolded protein stress response inducing brefeldin-a, dopamine receptor antagonist amisulpride, and pro-apoptosis *Bcl-2* inhibitor ABT-737 (Fig 4d). RNAi knockdown signatures with high connectivity to *Clic* knockdown included genes associated with cytoskeleton and membrane dynamics (*Josd1, Alms1, Tfg*), apoptosis (*Tnfaip3, Gsdmb, Tp53*), metabolism (*Pgm1, Acly, Etfb*), and translation (*Eif2s2*) among others (Fig 4d).

### Ethanol Sensitivity Altered by *Clic* Knockdown is Modulated by Hyperoxia

Considering the overrepresentation of differentially expressed genes related to oxidation-reduction processes (Fig 3), we investigated whether *Clic* knockdown flies may have a vulnerability or resistance to oxidative stress such as hyperoxia. However, under hyperoxic conditions, knockdown flies showed only a slight resistance, having a mean survival time of 175 hours compared to 171 hours for controls (S2 Fig, panel a). Considering that *Drosophila Clic* knockdown increases resistance to ethanol sedation (27), we explored possible effects of hyperoxia on ethanol sedation in *Clic* knockdown flies. As expected, knockdown of *Clic* blunted ethanol sedation sensitivity in flies housed under ambient (i.e. normoxia) conditions (Fig 5a-c, black bars). While exposure to hyperoxia for 1-3 days had no effect on ethanol sedation in a wild-type control strain (S2 Fig, panel b) or in *elav*/+ controls (Fig 5a-c), hyperoxia treatment significantly blunted—and in fact appeared to fully suppress—the ethanol sedation resistance observed in *Clic* knockdown flies under normoxia (Fig 5a-c, red bars). Furthermore, the blunting of resistance to ethanol sedation in the knockdown flies appeared to increase with the duration of hyperoxia exposure (Fig 5a-c). Interestingly, the v105975/+ genotype with limited knockdown of *Clic* exhibited an intermediate ethanol sedation resistance phenotype as previously reported (27), that was also suppressed by exposure of flies to hyperoxia (Fig 5a-c).

**Fig 5.**
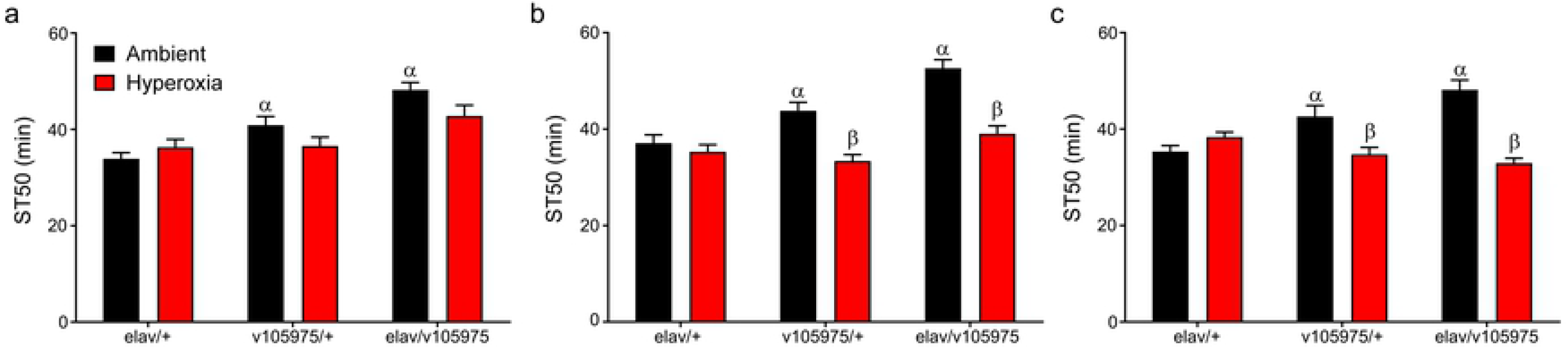
Ethanol Sensitivity Under Hyperoxia. Effect of chronic hyperoxia on acute ethanol sedation. ST50 is the time required for 50% of flies to become sedated. Longer ST50 represent resistance to ethanol sedation. (a) Day 1: Effect of Genotype (p<0.0001) but not hyperoxia (p=0.0950) and no interaction (p=0.0626). ^α^Effect of genotype under ambient conditions: ST50 longer in v105975/+ and *elav*/v105975 compared to control *elav*/+ (p<0.0001-0.0477). (b) Day 2: Effects of hyperoxia (p<0.0001) and genotype (p<0.0001) with a significant interaction (p=0.0021). ^α^Effect of genotype under ambient conditions: ST50 was longer in v105975/+ and *elav*/v105975 compared to control *elav*/+ (p<0.0001-0.0358). ^β^Within genotype, hyperoxia decreased ST50 (p<0.0001-0.0003). (c) Day 3: Effect of hyperoxia (p<0.0001) but not genotype (p<0.0791), and a significant interaction (p=0.0001). ^α^Effect of genotype under ambient conditions: ST50 was longer in v105975/+ and *elav*/v105975 compared to control *elav*/+ (p<0.0001-0.0172). ^β^Within genotype, hyperoxia decreased ST50 (p<0.0001-0.0078). Strain and hyperoxia conditions evaluated with two-way ANOVAs and Bonferroni’s multiple comparison post-tests.

## Discussion

The present study constitutes the first published transcriptomic profiling of a chloride intracellular channel genetic manipulation. We targeted *Clic*, the sole *Drosophila* chloride intracellular channel gene, for RNAi knockdown and performed differential gene expression and bioinformatic analysis to gain insight into the genes and biological processes perturbed by *Clic* reduction and to better understand the role of this gene in acute ethanol sedation sensitivity. Chloride intracellular channels are an enigmatic class of proteins, having characteristics of metamorphic proteins (7), ion channels (8), and redox enzymes (17). While previous studies have sought to identify chloride intracellular channel functions through more direct lines of investigation, such as *in vitro* assays of enzymatic reduction (17) and ion channel efflux capabilities (8), the present study has taken a more discovery-oriented approach by seeking to identify genes that respond to a reduction in *Clic* expression. Impressively, a neuronally-selective 41% knockdown of *Clic* altered the expression over 9% of the known *Drosophila* genome. Over-representation analysis of these differentially regulated genes identified several enriched GO terms including Oxidation-Reduction *Biological Process* and Membrane *Cellular Component* as well as significant overlap with gene sets from *Drosophila* ethanol sedation sensitivity and exposure studies. Extending our findings from *in silico* to *in vivo*, we evaluated *Clic* knockdown flies for sensitivity to ethanol sedation in the presence of hyperoxia and observed a blunting of sensitivity. Taken together, the studies published here provide additional evidence for known chloride intracellular channel functions and suggest that oxidative-reduction related gene expression may have an important role in *Clic* modulation of sensitivity to acute ethanol.

While inducible gene expression systems are invaluable for producing temporally and spatially precise genetic manipulations, they are often prone to leakage and the Gal4-UAS system is no exception. Leakage has previously been described for both Gal4 inducers and UAS transgenes, but extent of leakage is difficult to predict and can vary according to fly strain and age among other factors (51). Here we observe an intermediate phenotype in RNAi-only animals that fell between the knockdown and Gal4 strains in terms of gene expression and sensitivity to ethanol sedation. While the differential gene expression observed in the RNAi-only control was substantial, these are almost entirely the same set of genes differentially expressed in the Gal4-regulated knockdown strain. However, leaky expression could potentially complicate interpretation of the neuron-selectivity of the knockdown. Although the majority of the knockdown is occurring under the neuron-specific *elav*-Gal4 inducer, some component of the gene expression or ethanol sedation changes may be occurring in other cell types.

Overrepresentation analysis performed on *Clic* knockdown-responsive genes yielded multiple enriched GO terms of interest that both highlight known functions related to chloride intracellular channels but also point to possibly novel, undescribed roles. Chloride intracellular channels are known to interact with membranes, forming intracellular channels (7,52), associating with membrane domains undergoing tubulogenesis (2,53), and promoting membrane trafficking (9,10). These activities correspond well to the GO term hits, Lipid Particle and Membrane. Furthermore, CMap analysis identified knockdown of *Josd1, Alms1*, and *Tfg*, three genes with functions linked to cytoskeleton and membrane dynamics, as being highly connectivity to the *Clic* knockdown signature. A similar GO term hit, Cell Junctions, has relevance to vertebrate *Clic* orthologs, which have been shown to be enriched at junctions between dividing cells, where they are potentially regulating cytoskeletal organization (54).

The GO term Oxidation-Reduction Process was enriched in *Clic* knockdown-sensitive genes and may reflect a known role of chloride intracellular channels in carrying out oxidoreductase reactions (17). Although evidence for this function is limited to observation *in vitro*, it has been long suspected based on the homologous omega class glutathione S-transferase structure of chloride intracellular channels (1,6). Thus, our transcriptome analysis validates the prior in vitro studies on a role of *Clic* in oxidation-reduction. Also supporting known roles for chloride intracellular channels, *Clic* knockdown showed high connectivity on CMap analysis with the apoptosis-blocking drug ABT-737 and with pro-apoptosis gene p53. It has been shown that chloride intracellular channels have a p53 binding element in its promoter, upregulate in response to various cell stressors including DNA damage, and has been shown to traffic to the nucleus as an early responder to cell stress where it also participates in apoptosis (11,12). A potentially novel association of *Clic* identified in this study is protein translation, for which Cytoplasmic Translation was the top GO term from the overrepresentation analysis and was enriched almost exclusively by downregulated genes. In concordance with this, CMap analysis showed a strong negative connectivity between the *Clic* knockdown signature and translation initiation factor, *Eif2s2*. Also potentially novel, CMap analysis identified multiple histone deacetylase inhibitors with strong connectivity to *Clic* knockdown.

Chloride intracellular channels are highly conserved evolutionarily and vertebrates possess a family of 6 paralogs (Littler 2010). *Drosophila Clic* has high sequence similarity to vertebrate orthologs including *Clic4*, which has been shown to be regulated by ethanol (25,26) and capable of decreasing ethanol sedation sensitivity when overexpressed in mouse brain (25). Neuronal *Drosophila Clic* knockdown has previously been shown to decrease ethanol sedation sensitivity (27), consistent with our findings here, showing a conservation of function between mouse and *Drosophila* orthologs. Of note, the decreased sensitivity to ethanol sedation is obtained through opposing genetic manipulations in mice and flies, overexpression and knockdown, respectively. As hypothesized previously, this difference in phenotype expression may be due to species-specific differences in number and presence of chloride intracellular channel paralogs or the experimentally targeted cell types or brain regions (25). Novel to this body of work, we show that while *Clic* knockdown decreased sensitivity to ethanol sedation, this effect was reversed by hyperoxia in a time-dependent manner. Considering hyperoxia had no effect on the control strain, this decrease in ethanol sedation sensitivity with time in the knockdown strain suggests that biological functions altered by *Clic* knockdown, which decreases sensitivity to ethanol sedation, are also either regulated on some level by hyperoxia or interact functionally with molecular responses to hyperoxia. This possibility is underscored by overrepresentation of genes related to GO oxidation-reduction processes in both the *Clic* knockdown-responsive gene set and GeneWeaver overlap analysis with ethanol-related *Drosophila* gene sets. Furthermore, metabolism of ethanol produces reactive oxygen species and cellular oxidative stress while oxidoreductase enzymatic activity has been reported of vertebrate chloride intracellular channels *in vitro* (17). The exact molecular interactions between *Clic*, ethanol and hyperoxia thus merit future investigation.

Remarkably, nearly one third of genes responsive to *Clic* knockdown were found to be shared with a union set of published ethanol sedation sensitivity-related *Drosophila* genes. Three of these gene sets display ethanol regulation during acute exposure (44–46) while the fourth represents genes differentially expressed between strains artificially selected for high and low ethanol sedation sensitivity (47). This intersection between *Clic* knockdown-responsive and ethanol-regulated genes suggests a major role for *Clic* in molecular pathways governing ethanol sedation sensitivity and the acute response to ethanol. Functional enrichment of the shared gene set implicates a variety of possible processes including amino acid metabolism, oxidation-reduction, sensory perception, protein processing, and transport.

Employing the Gal4-UAS system, this study is the first to characterize the transcriptome following genetic manipulation of a chloride intracellular channel gene. Bioinformatic analysis of knockdown-induced differentially regulated genes provided support for existing evidence that *Clic* is involved in oxidation and reduction processes and has roles near cellular membranes. Novel to this work, we also identified an enrichment of *Clic* knockdown-sensitive genes related to cytoplasmic translation and heme binding and associated with the nucleolus and cell junction. We have also determined that an interaction between hyperoxia and *Clic* expression modulates ethanol sedation sensitivity. Taken together, these studies add to the growing body of literature supporting *Clic* genes as important for ethanol-related behaviors and also being involved in redox-related processes.

## Acknowledgments

The authors thank Brandon Shell for technical assistance and the Vienna *Drosophila* RNAi Center and the Bloomington *Drosophila* Stock Center for *Drosophila* strains. We also acknowledge the excellent assistance of Tana Blevins of the VCU Molecular Diagnostics Core Laboratory for processing of microarrays.

## Supporting Information

**S1 Fig: Differential Gene Expression by Strain**. (a) Differentially regulated genes (FDR < 0.05) for each possible fly strain contrast. (b) Genes differentially expressed between knockdown (*elav*/v105975) and RNAi-only control (v105975) are also altered in the knockdown vs Gal4-only control (*elav*/+) contrast.

**S2 Fig: Hyperoxia Survival and Control Strain Sedation Sensitivity**. (a) Survival analysis for flies exposed to continuous hyperoxia grouped by strain. (b) Ethanol sedation times for wild-type control flies under ambient and hyperoxic conditions for 3 days.

**S1 Table: Differentially Expressed Genes**

**S2 Table: Enriched Gene Ontology Terms**

**S3 Table: GeneWeaver Ethanol Gene Sets**

